# Combined Right Ventricular Preload and Afterload Challenge Induces Contractile Dysfunction and Ventricular Arrhythmia in an Age-dependent Manner in Isolated Hearts of Dystrophin-deficient Mice

**DOI:** 10.1101/2023.11.14.564005

**Authors:** Andrew Behrmann, Jessica Cayton, Matt Hayden, Michelle Lambert, Zahra Nourian, Keith Nyanyo, Laurin Hanft, Maike Krenz, Kerry McDonald, Timothy Domeier

## Abstract

Duchenne muscular dystrophy (DMD) is an X-linked recessive myopathy due to mutations in the dystrophin gene. Diaphragmatic weakness in DMD causes restrictive lung disease and excessive afterload on the right ventricle (RV). Thus, RV dysfunction in DMD develops early in disease progression. Herein, we deliver a 30-minute sustained RV preload and afterload challenge to isolated hearts of wildtype (Wt) and dystrophic (mdx^4CV^) mice at both young (2-6 month) and middle-age (8-12 month). Young dystrophic hearts exhibited greater pressure development than wildtype under unloaded conditions but following RV preload/afterload challenge exhibited similar contractile function as wildtype. Following RV preload/afterload challenge, young dystrophic hearts had an increased incidence of ventricular arrhythmias compared to wildtype. Hearts of middle-aged wildtype and dystrophic mice had similar contractile function during unloaded conditions, yet, after RV preload/afterload challenge, hearts of middle-aged dystrophic mice had severe RV contractile dysfunction and ventricular arrhythmogenesis, including ventricular tachycardia. Following load challenge, young and middle-aged dystrophic hearts had greater damage, measured via tissue LDH release, than wildtype mice of similar age. Our data indicate complex age-dependent changes in RV function and arrhythmogenesis with load in dystrophin deficiency, highlighting the need to avoid sustained RV load to forestall RV dysfunction and arrhythmia.

## Introduction

Duchenne muscular dystrophy (DMD) is a progressive and degenerative myopathy that occurs due to mutations in the *DMD* gene on the X-chromosome encoding the cytoskeletal protein dystrophin (1, 2). Dystrophin is the main cytoplasmic component of the dystrophin-glycoprotein complex that is present in all striated muscle fibers and plays a key role in maintaining the integrity of the sarcolemma by linking it to the actin cytoskeleton (3, 4). Skeletal muscle lacking dystrophin is highly susceptible to contraction-induced sarcolemmal damage and cellular necrosis (5, 6). Cardiomyocytes lacking dystrophin have also shown sarcolemmal damage with passive distention, which led to a stiffened, hypercontractile state and eventual cell death due unregulated influx of calcium and fibrotic damage (7, 8).

Clinically, DMD is the most common muscular dystrophy, effecting 19.8 per 100,000 live male births, and a pooled global prevalence of 7.1 per 100, 000 males (9). Patients with DMD experience progressive skeletal muscle wasting and diaphragmatic weakness leading to severe respiratory complications. Eventually, DMD patients lose almost all ambulatory function and have a shortened life expectancy of 30 to 40 years with the main cause of death being cardiopulmonary failure (10). It has been well documented that respiratory function, pulmonary vascular resistance (right ventricular afterload), and right ventricular function are tightly coupled and that elevated pulmonary vascular resistance can lead to deleterious effects on the right ventricle (11, 12). Thus, it has been noted that the diaphragm weakness in DMD leads to pulmonary hypertension (increased right ventricular afterload) and right ventricular failure (13). Patients suffering from DMD are also at increased risk of suffering from ventricular arrhythmia (14). It is crucial to gain a better understanding of both the mechanisms and triggers of RV dysfunction and ventricular arrhythmia in dystrophic patients to improve outcomes.

The dystrophin-deficient mdx mouse displays the most severe fibrosis within the diaphragm, making it a clinically relevant model to study the intricate coupling of cardiopulmonary function (15, 16). This is exemplified by the fact that restoring only diaphragm muscle functionality to dystrophic mice is cardioprotective and can restore both right and left ventricular ejection fractions (17). Right ventricular dysfunction has been shown to precede left ventricular dysfunction in dystrophic mice starting at 3 months of age showing an increased end systolic volume and reduced RV ejection fraction (18). Furthermore, right ventricular fibrosis has been shown to precede fibrosis of the left ventricle and interventricular septum. Additionally, an increase in preload via abdominal compression significantly altered LV systolic pressures; however, there was no significant difference between dystrophic and wildtype control hearts in RV systolic pressure when preload was increased (19). Despite the RV preload alterations in the aforementioned study, right ventricular afterload was not altered nor controlled despite it being a key feature of right ventricular failure in both DMD and pulmonary hypertension. This necessitates additional investigations into right ventricular function not only in a dystrophic mouse model, but for other rodent models that investigate the interplay of the cardiopulmonary system and right ventricular function.

Hence, the purpose of the current study is to extend previous studies in dystrophic mouse models by assessing the effects a sustained and controlled elevation in right ventricular load in ex vivo hearts and examine the severity of right ventricular phenotypes of young and middle-aged dystrophic mice compared to wildtype mice of the same age to provide insight on right ventricular function with aging in DMD.

## Methods

### Animal model

All protocols in this paper involving animals were performed in accordance with the Animal Care and Use Committee (Approval reference number 35701) of the University of Missouri. This study used wildtype C57Bl/6 mice (Wt) and dystrophin-deficient B6Ros.Cg- *Dmd^mdx-4Cv^*/J (Jackson labs strain #002378), (Dmd^mdx-4Cv^) male mice that were 2-6 months (young) or 8-12 months (middle-aged) of age.

### Solutions

A modified Krebs-Henseleit buffer (KHB) was used for working heart perfusion. The KHB contained the following (in mmol/L): 117 NaCl, 4.7 KCl, 1.2 MgSO4, 1.2 KH2PO4, 25 NaHCO3, 11.1 glucose, 0.4 caprylic acid, 1 pyruvate, 0.0023Na EDTA, and 1.8 CaCl2. Solution was warmed to 37℃ in water-jacketed glassware and maintained via a recirculating pump (Radnoti, #170051G).

### Isolated Working Heart

Mice were injected with intraperitoneal ketamine:xylazine (100 mg/kg:5mg/kg), and once there was an absence of the pedal withdrawal reflex, hearts were rapidly excised within 30 seconds (20). Once excised, hearts were immediately placed in cold (4℃) KHB without calcium. The hearts were cleared of excess lung, adipose, and aortic tissue then submerged in a cold KHB bath also lacking calcium and then cannulated via the aorta (60 mmHg afterload) to deliver warm (37℃) oxygenated KHB with 1.8 mM calcium to perfuse the coronary circulation (Langendorff mode). Any blood and cold KHB within the bath from the initial cannulation were suctioned out of the bath and replaced with warm oxygenated KHB containing 1.8 mM calcium before placing additional cannulas. The right atrium and the pulmonary artery were subsequently cannulated. Perfusate flow into the right atrium was closed via a stopcock during unloaded baseline conditions and opened to a preload of 10 mmHg for preload challenge. The pulmonary artery cannula was connected to a fluid column with a three-way stopcock set at the level of the heart that was either open to atmosphere (unloaded conditions) or connected to a 20 mmHg afterload column for the afterload challenge. An 22-gauge needle was used to make a small insertion hole in the apical portion of the right ventricle and a 1.0 F Millar pressure catheter was inserted to monitor right ventricular pressures using an FE231 Bio Amp and LabChart/Power 8.1 software (AD Instruments) (Figure1A). The sampling rate was set to 1000 Hz with a 10 Hz low pass filter applied to improve signal-to-noise of pressure waveforms. After a 15-minute unloaded baseline period in Langendorff mode, the right ventricle was then challenged with 10 mmHg preload and 20 mmHg afterload for 30 minutes. The three 5-minute intervals that right ventricular function and arrhythmias were assessed were: 1) final 5 minutes of baseline Langendorff mode conditions, 2) initial preload/afterload challenge (maximum response to pressure elevation), and 3) 30 minutes post right ventricular stretch (Figure 1B). All measurements were gathered as an average of 5-10 seconds of steady state pressure and ECG tracings.

**Figure 1:**
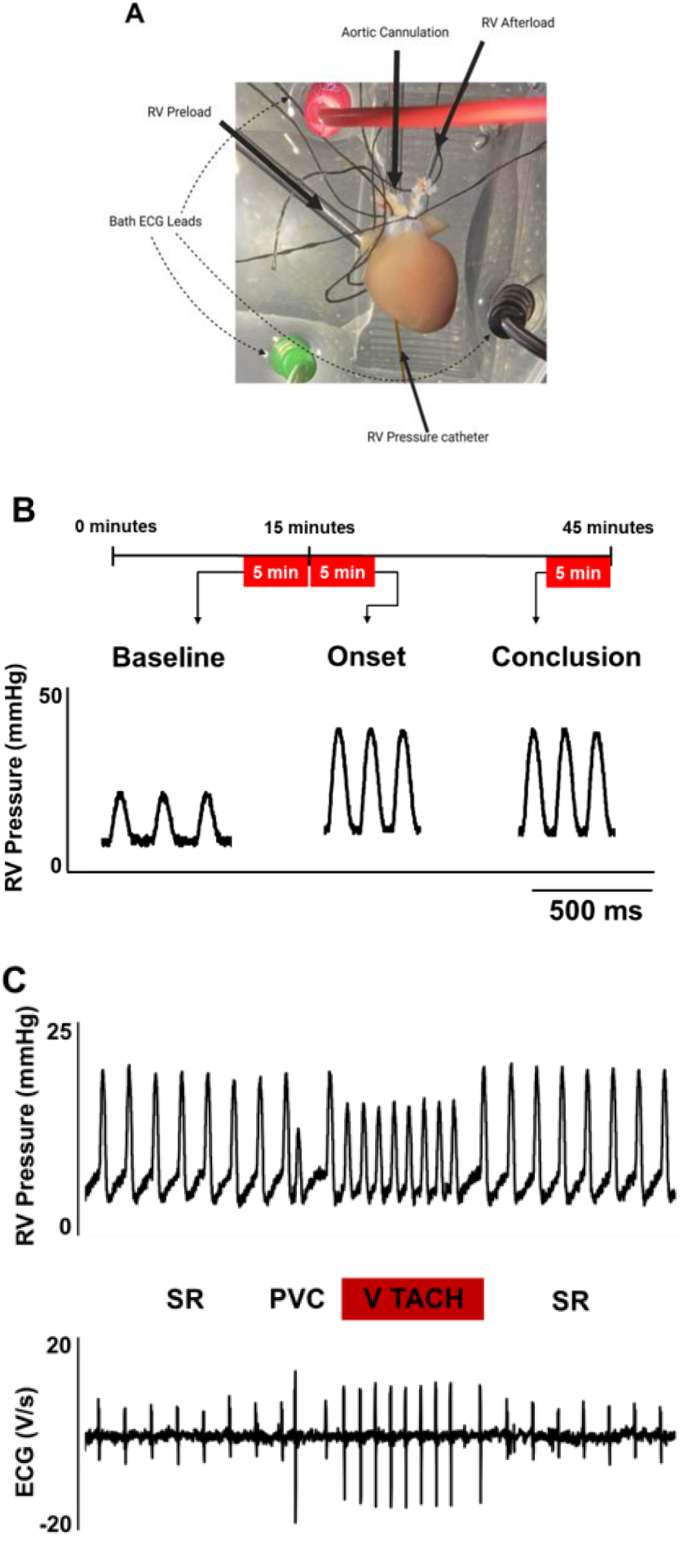
Cannulations with pressurization protocol and example tracings. (**A**) Isolated heart preparation and (**B**) preload/afterload challenge protocol with representative right ventricular pressure tracings. (**C**) Example pressure traces (upper) and ECG traces (lower) indicating sinus rhythm (SR), a premature ventricular contraction (PVC), and short bout of ventricular tachycardia (V TACH).

### Electrocardiogram (ECG) Measurements

Arrhythmias were assessed at baseline and during the right ventricular load challenge using a set of three 1.5 mm shrouded socket monopolar 29-gauge MLA1213 needle electrodes connected to a FE231 Bio Amp and LabChart/Power 8.1 (AD Instruments). Electrode position was adjusted as needed to gain clear distinction of each cardiac cycle’s P wave and QRS complex. A premature ventricular contraction (PVC) was defined as a premature ventricular complex on the ECG waveform that either preceded or was completely independent of an atrial driven P wave. The QRS complexes of PVCs tended to be of higher amplitude and longer duration compared to the QRS complexes of sinus rhythm, allowing for clear identification. The incidence of ventricular arrhythmia within each minute of the five-minute intervals was quantified using a 0-1-2 scoring system. A score of 0 indicating no arrhythmias within that minute, a score of 1 indicating isolated PVCs, and score of 2 indicating the presence of salvos of 2-3 PVCs or runs of ventricular tachycardia/fibrillation (Figure 1C) (21). The five-minute average arrhythmia score for each heart was recorded for further data analysis and comparison.

### LDH Effluent Collection

Following the right ventricular load challenge, coronary effluent was collected for detection and quantification of lactate dehydrogenase (LDH) using the Promega luminescent LDH-Glo 105TM Cytotoxicity Assay kit following manufacturer protocol (Promega, Madison, WI, USA).

### Statistical Analysis

All samples are reported as mean ± standard error of the mean. To compare group means, a two-sided student’s t-test was performed in Graph Pad/Prism. Analysis of LDH bioluminescent assay was performed using a one-tailed Mann-Whitney U-Test. A p-value <0.05 was considered statistically significant (22).

## Results

### Young

Compared to wildtype mice under unloaded conditions, young dystrophic hearts exhibited greater right ventricular Pdev (Figure 2A). Both young groups exhibited similar Pdev at the onset (Figure2B) and conclusion (Figure2C) of the combined preload/afterload challenge. Change in pressure development from the onset to the conclusion of the preload/afterload challenge did not significantly differ between young wildtype and young dystrophic mice (Figure2D). Average heart rates did not significantly differ between young wildtype and dystrophic hearts during unloaded conditions (Heart rate: 368±29 wildtype versus 355±45 mdx^4Cv^, P=0.56), initial loaded conditions (Heart rate: 433±24 wildtype versus 447±18 mdx^4Cv^, P=0.14), and the conclusion of loaded conditions (Heart rate: 401±29 wildtype versus 426±22 mdx^4Cv^, P=0.16). Arrhythmia incidence was low in young wildtype and dystrophic hearts during unloaded conditions (Figure 3A) yet increased in dystrophic hearts at both the onset (Figure 3B) and conclusion of the preload/afterload challenge (Figure 3C). There were no bouts of ventricular tachycardia observed in young wildtype or dystrophic mice, only isolated PVCs. LDH release following the preload/afterload challenge was significantly higher in young dystrophic hearts compared to young wildtype hearts (Figure 4).

**Figure 2:**
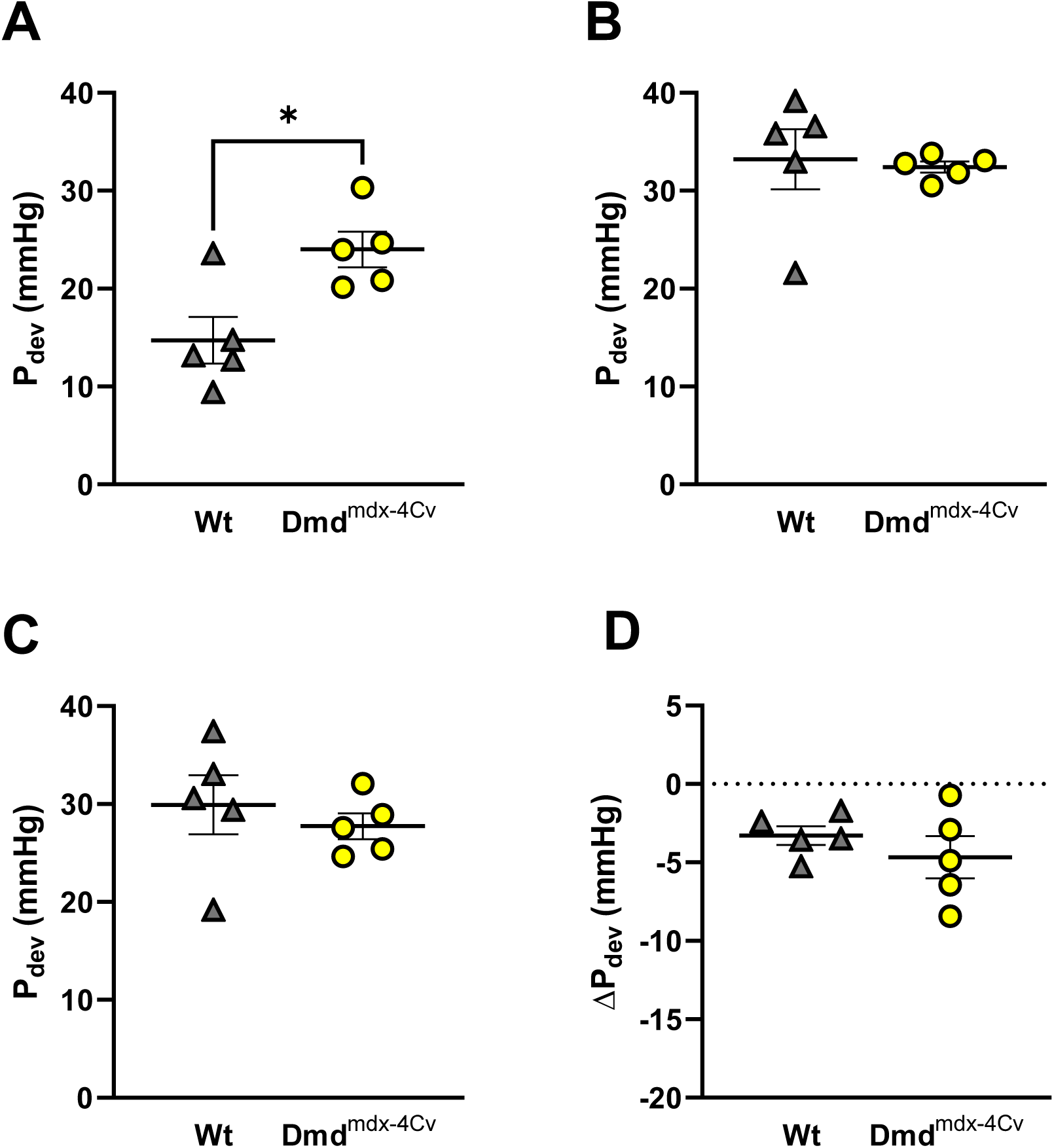
Right Ventricular Pressure Development of Young wildtype and dystrophic Dmd^mdx-4Cv^ hearts. Summary data of Developed Pressure (Pdev) of Young Wt (n=5, grey triangles) and Dmd^mdx-4Cv^ (n=5, yellow circles) at last five minutes of baseline (**A**), first five minutes of preload/afterload challenge (**B**), and last five minutes of preload/afterload challenge (**C**). Change in Pdev from the first five minutes to the last five minutes of the preload/afterload challenge shown in (**D**). * p = 0.015 in (**A**), Independent-samples t-test.

**Figure 3:**
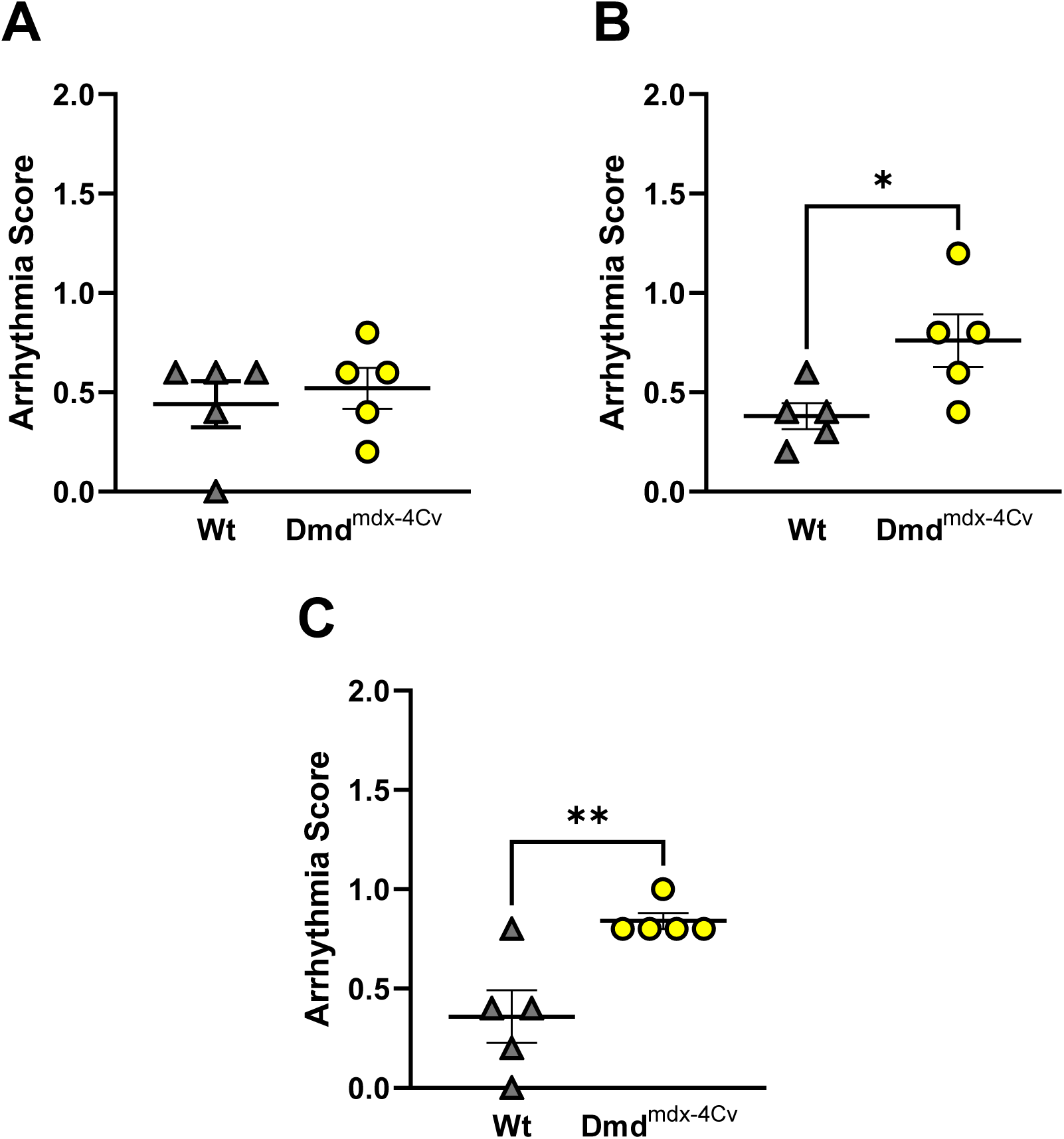
Ventricular Arrhythmia incidence in Young wildtype and dystrophic Dmd^mdx-4Cv^ hearts. Summary data of average arrhythmia scores of Young Wt (n=5, grey triangles) and Dmd^mdx-4Cv^ (n=5, yellow circles) at last five minutes of baseline (**A**), first five minutes of preload/afterload challenge (**B**), and last five minutes of preload/afterload challenge (**C**). * p = 0.03 in (**B**), ** p = 0.009 in (**C**), Independent-samples t-test.

**Figure 4:**
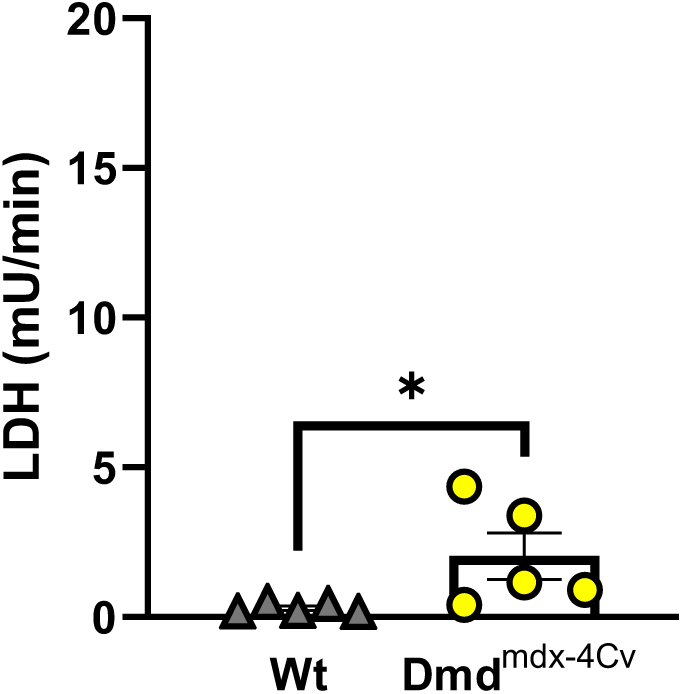
Cardiac lactate dehydrogenase release following preload/afterload challenge in Young wildtype and dystrophic Dmd^mdx-4Cv^ hearts. Summary data of lactate dehydrogenase (LDH) release of Young Wt (n=5, grey triangles) and Dmd^mdx-4Cv^ (n=5, yellow circles) hearts at the conclusion of the preload/afterload challenge. * p = 0.016, one-tailed Mann-Whitney U-test.

### Middle-aged

In middle-aged mice, hearts of both wildtype and dystrophic groups exhibited similar Pdev under unloaded conditions (Figure 5A). However, Pdev was lower in dystrophic hearts at both the onset (Figure 5B) and conclusion (Figure 5C) of the preload/afterload challenge.

**Figure 5:**
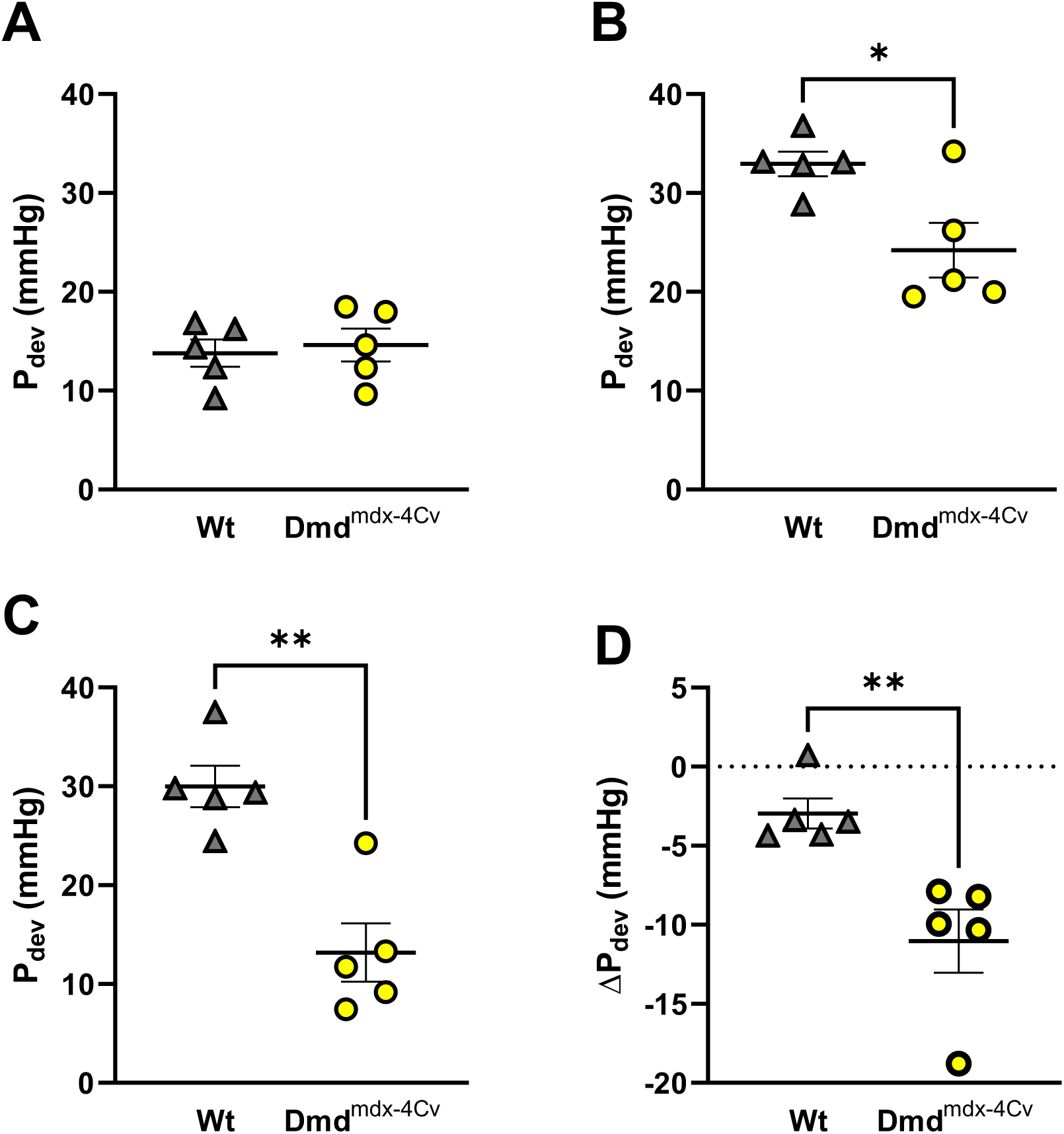
Right Ventricular Pressure Development of Middle-age wildtype and dystrophic Dmd^mdx-4Cv^ hearts. Summary data of Developed Pressure (Pdev) of Middle-age Wt (n=5, grey triangles) and Dmd^mdx-4Cv^ (n=5, yellow circles) at last five minutes of baseline (**A**), first five minutes of preload/afterload challenge (**B**), and last five minutes of preload/afterload challenge (**C**). Change in Pdev from the first five minutes to the last five minutes of the preload/afterload challenge shown in (**D**). * p = 0.017 in (**A**), ** p = 0.002 in (**C**), ** p = 0.006 in (**D**), Independent-samples t-test.

Furthermore, during the sustained load challenge, middle-aged dystrophic hearts exhibited a significant loss of right ventricular contractile function (Figure 5D). Average heart rate did not significantly differ between middle-aged wildtype and dystrophic hearts during unloaded conditions (Heart rate: 386±25 wildtype versus 384±19 mdx^4Cv^, P=0.89), initial loaded conditions (Heart rate: 410±15 wildtype versus 409±18 mdx^4Cv^, P=0.19), and the conclusion of loaded conditions (Heart rate: 423±18 wildtype versus 425±24 mdx^4Cv^, P=0.88). Arrhythmia incidence was low in middle-aged hearts of both groups during unloaded conditions (Figure 6A). However, at the onset (Figure 6B) and conclusion (Figure 6C) of the combined preload/afterload challenge, middle-aged dystrophic hearts exhibited significantly higher arrhythmia scores. Bouts of ventricular tachycardia were only observed in middle-aged dystrophic hearts (n=4/5 hearts). Middle-aged dystrophic hearts also had significantly greater LDH release compared to middle-aged wildtype hearts at the conclusion of the preload/afterload protocol (Figure 7).

**Figure 6:**
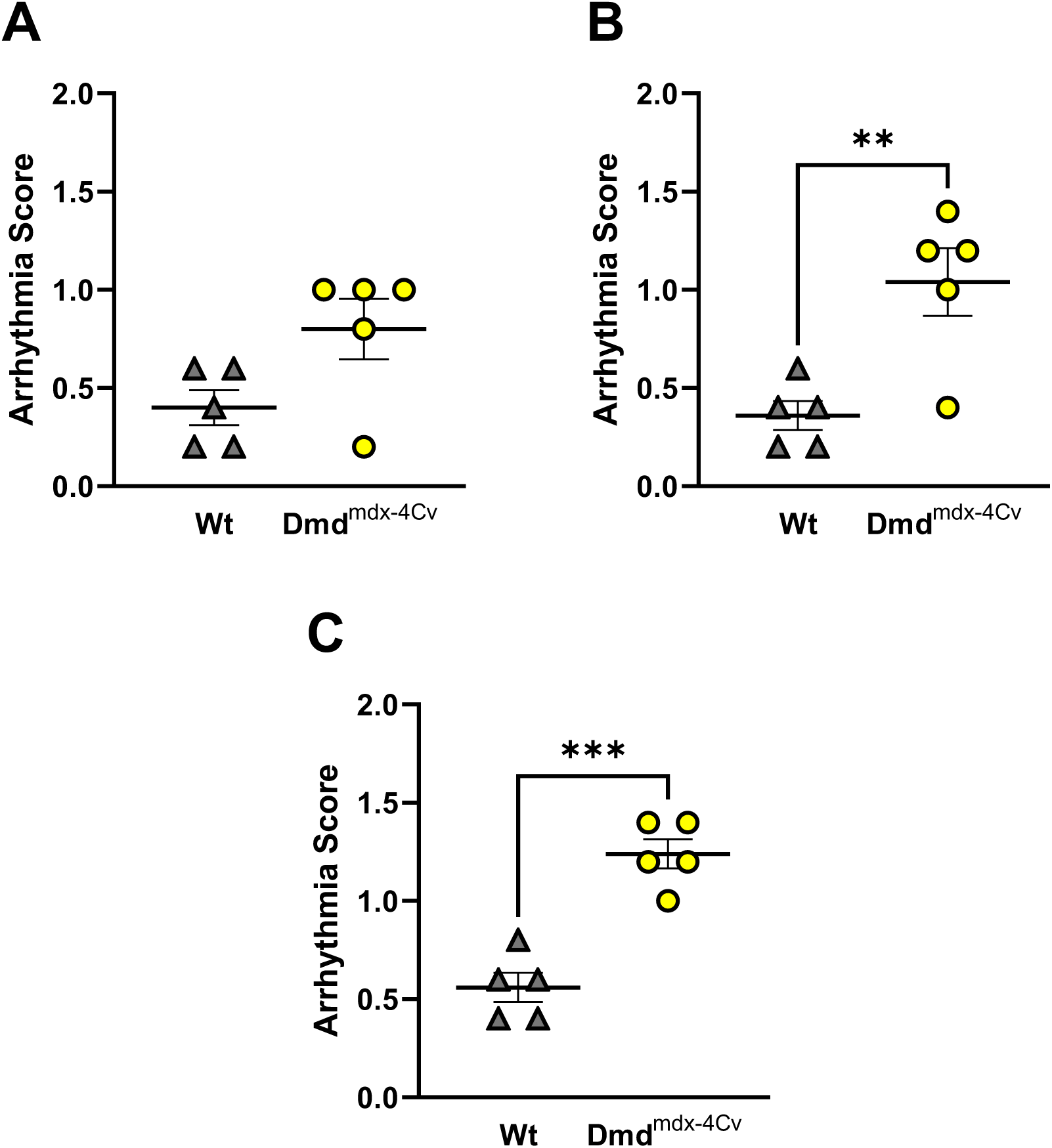
Ventricular Arrhythmia incidence in Middle-age wildtype and dystrophic Dmd^mdx-4Cv^ hearts. Summary data of average arrhythmia scores of Middle-age Wt (n=5, grey triangles) and Dmd^mdx-4Cv^ (n=5, yellow circles) at last five minutes of baseline (**A**), first five minutes of preload/afterload challenge (**B**), and last five minutes of preload/afterload challenge (**C**). ** p = 0.0067 in (**B**), *** p = 0.0007 in (**C**), Independent-samples t-test.

**Figure 7:**
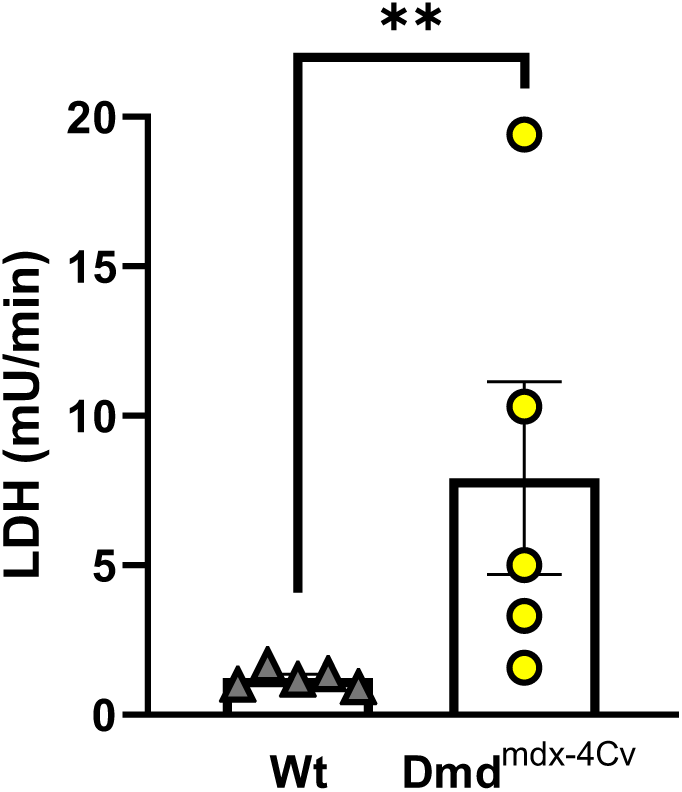
Cardiac lactate dehydrogenase release following preload/afterload challenge in Middle-age wildtype and dystrophic Dmd^mdx-4Cv^ hearts. Summary data of lactate dehydrogenase (LDH) release of Middle-age Wt (n=5, grey triangles) and Dmd^mdx-4Cv^ (n=5, yellow circles) hearts at the conclusion of the preload/afterload challenge. ** p = 0.008, one-tailed Mann-Whitney U-test.

## Discussion

This study examined right ventricular function in a mdx^4Cv^ mouse model using an isolated working heart method aimed at fundamentally mimicking the RV disease phenotype previously described in dystrophic mice and patients (13, 19, 23). In vivo MRI of mdx mice noted that right ventricular dysfunction is seen as early as 3 months, showing elevated end systolic volume and a decreased ejection fraction (24). Notably, our study found that during unloaded baseline conditions, young dystrophic hearts were hypercontractile compared to wildtype, but middle-aged hearts displayed similar contractility while unloaded. This discrepancy may be explained by the fact that under baseline conditions, the isolated mdx heart may be in a hypercontractile state to compensate for the elevated afterload seen in vivo (25). However, our data suggest that dystrophic hearts lose an adaptive hypercontractile mechanism with aging, and exhibit a concomitant loss in the Frank-Starling response (Figure 5B). Furthermore, the middle-aged dystrophic hearts exhibited a remarkable loss in right ventricular contractile function with sustained elevation in right ventricular load (Figure 5C,D). The elevated RV afterload seen in DMD-induced pulmonary hypertension can result in cardiac afterload stress and damage (7, 21, 26), which can lead to cellular hypercontractility and eventual cell death. Our study found similar results with dystrophic hearts releasing significantly more LDH compared to wildtype hearts of the same age regardless of maintained (young) or diminished (middle-age) cardiac function throughout the load challenge. Presumably, further disease progression will result in the damaged tissue being replaced by fibrous tissue which, in turn, further worsens overall RV function.

The dystrophic mouse models have previously shown susceptibility to ventricular arrhythmias while using programmed electrical stimulation and a catecholamine challenge (27, 28). Similarly, in vivo ECG recordings revealed there was an increased incidence of PVCs in dystrophic versus wildtype mice which could trigger persistent ventricular tachycardias (29). Indeed, our group recently determined that increased preload in the left ventricle of mdx^4Cv^ hearts led to calcium handling abnormalities and increased incidence of ventricular arrhythmias (21). The current study found that dystrophic mouse hearts under elevated right ventricular load also had an increased incidence of arrhythmia, however the incidence and severity of arrhythmia increased with age (c.f., Figure 3 and Figure 6) consistent with adverse ventricular remodeling at a later stage of disease. The increased incidence of ventricular tachycardia in middle-aged dystrophic mice during the combined preload/afterload challenge highlights the need to avoid sustained periods of elevated right ventricular load in dystrophic patients (e.g. suboptimal ventilatory support) as this may increase the risk of ventricular arrhythmia and sudden cardiac death.

In conclusion, our study suggests that unloaded right ventricular hypercontractility displayed in young dystrophic hearts may be a temporary, adaptive mechanism to compensate for the pulmonary hypertension observed in dystrophic mice (19, 30, 31). As baseline hypercontractility disappears in middle-age dystrophic hearts, the ability to withstand prolonged elevated loading conditions also declines. An inability to meet afterload demands in vivo may lead to increased end systolic volumes and thus, preload elevation (24, 30). Furthermore, there is a plethora of data that describes the right ventricle of dystrophic mice displays severe fibrosis that worsens with age (19, 24, 25, 32, 33). The fibrosis may serve as a substrate for ventricular arrhythmia upon stretch-induced calcium mishandling, thus acting as the triggering event that increases arrhythmia incidence and severity in middle-aged dystrophic hearts compared to wildtype mice of similar age (21, 34). Clinically, these data highlight that DMD patients should avoid prolonged situations of excessive RV load due to risk of fatal ventricular arrhythmia.

## Notes

### Competing Interest Statement

The authors have declared no competing interest.

